# Distinct Response Patterns to PHB Modulators in B Cell Lymphoma Models

**DOI:** 10.1101/2025.06.12.659346

**Authors:** Christoph Schultheiß, Maja Kadel, Laurent Désaubry, Mascha Binder

## Abstract

B cell malignancies, including chronic lymphocytic leukemia (CLL) and diffuse large B cell lymphoma (DLBCL), rely on dysregulated B cell receptor (BCR) signaling for survival and proliferation. Prohibitin 1 and 2 (PHB1, PHB2) are multifunctional proteins involved in mitochondrial function, IgM-type BCR signaling and other key oncogenic pathways, making them potential therapeutic targets in lymphomas. Here, we assessed the effects of five PHB-targeting small molecules - FL3, Mel6, Mel56, IN44, and Fluorizoline - on lymphoma cell lines as proof-of-concept study. PHB transcript and protein quantities were differentially affected and distinct patterns of antiproliferative effects and viability were observed. Across cell models, FL3, Mel56, and Mel6 displayed strongest effects. FL3 and Mel56 exerted strong cytotoxic effects, while Mel6 primarily slowed proliferation. IN44 showed modest but selective cytotoxic effects in an ABC-DLBCL model, while Fluorizoline selectively stopped proliferation of a Burkitt lymphoma model. Non-malignant stromal cells remained largely unaffected by Mel56, highlighting a potential therapeutic window of this inhibitor. Replacing the native IgM constant region by IgG in the MEC-1 CLL line using CRISPR-Cas9 resulted in a somewhat reduced, but not abrogated effect of Mel56 suggesting effects on additional pathways beyond the BCR. Together these data provide proof-of-concept evidence for PHB inhibition as a potential strategy to target B cell lymphomas.

## Introduction

B cell malignancies represent a heterogeneous group of lymphoproliferative disorders marked by the aberrant survival and expansion of B lymphocytes^1-4^. A central feature of many B cell malignancies is their dependence on B cell receptor (BCR) signaling, which governs key processes such as proliferation, apoptosis resistance, and metabolic adaptation^1,4^. Aberrant activation of BCR signaling networks including downstream pathways like NF-κB, PI3K-Akt, and MAPK/ERK sustains disease progression and therapy resistance in various B cell neoplasms^5,6^. Thus, dissecting how BCR signaling is modulated across different disease contexts may uncover new promising avenues for more precise and effective therapeutic intervention^7^. The pleiotropic scaffold proteins Prohibitins 1 and 2 (PHB1, PHB2) may represent emerging targets in this regard. Beyond their roles in mitochondrial maintenance, apoptosis, and MAPK/Akt signaling, PHBs also contribute to immune signaling. PHB1 scaffolds Raf-1 to support MEK–ERK activation^8^, while PHB2 engages autophagy machinery in mitophagy and interacts with transcriptional regulators of immune responses^9-12^. In previous work on chronic lymphocytic leukemia (CLL), we identified PHB2 as a shared intracellular binding partner of both the IgM tail of the BCR and the cytoplasmic domains of SLAMF1 and SLAMF7^13^. High SLAMF receptor expression impairs BCR signal transduction by sequestering PHB2, ultimately influencing the responsiveness of CLL cells to BCR pathway inhibitors and modulating NK cell education within the CLL microenvironment^13^. These findings highlight a broader regulatory role of PHB2 in orchestrating BCR-related signaling networks in CLL.

In light of this data, therapeutic interference with these BCR modulators appears to be a promising option not only in CLL but across a spectrum of B cell malignancies. The pleiotropic functions of PHB proteins across cellular compartments position them as promising candidates for broader therapeutic intervention beyond conventional BCR-targeting approaches. Here, we present proof-of-concept data demonstrating that pharmacologic modulation of PHBs selectively impairs survival and proliferation in various B cell malignancies while sparing non-malignant cells. Using five distinct PHB-targeting small molecules, we explore the therapeutic vulnerability of malignant B cells to PHB interference, suggesting that PHB-directed strategies may offer a novel, broadly applicable treatment avenue in lymphoid cancers.

## Material and methods

### RNAseq data

Publicly available RNAseq data sets were accessed via the EMBL-EBI Expression Atlas^14^, cBioportal^15^, UCSC Xena^16^, and GEPIA2^17^. Data were plotted with GraphPad Prism 10 (GraphPad Software, La Jolla, CA, USA) or via web-implemented tools in cBioportal and GEPIA2. GEPIA2 accesses RNAseq data of the Cancer Genome Atlas (TCGA)^18^, RNAseq data for the cell models was derived from Klijn et al.^19^, cBioportal RNAseq data for DLBCL, Burkitt lymphoma was generated by Thomas et al.^20^ and by Knisbacher et al.^21^ for CLL.

### Cell models and propagation

The cell lines MEC-1, Raji, OCI-Ly1, OCI-Ly3, RI-1, DG-75 and HH were obtained from the German Collection of Microorganisms and Cell Cultures GmbH (DSMZ, Braunschweig, Germany), M2-10B4 cells from ATCC. MEC-1 cells were maintained in IMDM medium supplemented with 10% (v/v) fetal bovine serum and 1% penicillin/streptomycin. The remaining cells were cultured in RPMI 1640 with 10% (v/v) fetal calf serum (Raji, RI-1, DG-75, HH) or 20% FCS (OCI-Ly1, OCI-Ly3, OCI-Ly8) and 1% (v/v) penicillin/streptomycin. All cell lines were incubated at 37°C and 5% CO_2_.

### Cell line engineering

Depletion of endogenous SLAMF1/SLAMF7 was performed via CRISPR-Cas9 as described^13^. For re-expressing SLAMF1 or SLAMF7 in SLAMF1/7-KO cells, cDNAs were cloned into the pLeGO-iC2-Puro lentiviral vector. Lentiviral particles were generated by transient transfection of HEK293T cells using the third generation packaging plasmids pMDLg/pRR3 and pRSV-Rev and the VSV-G envelope protein^22^. Transduction was performed in the presence of polybrene (8 μg/ml) while centrifuging 1 hour at 1000*g* followed by selection with puromycin (1 µg/ml) as described^13,23^. The switched IgG MEC-1 model was generated using CRISPR-Cas9 as described^24^.

### Proliferation assays

To monitor proliferation, cells were seeded at 1 × 10^6^ cells / 10 ml in 25 cm^2^ flasks and treated with FL3 (50 nM), Mel6 (10 µM), Mel56 (10 µM), IN44 (10 µM) or Fluorizoline (1 µM). All compounds were dissolved in DMSO. Cells left untreated or treated with DMSO (10 µM) served as control. Every 24 hours, viable cells were quantified by trypan blue staining using the Cell Viability Analyzer Vi-Cell XR (Beckman Coulter). Heatmaps of proliferation and viability measures were generated with the R package pheatmap using the R version 4.3.1 and RStudio 2023.06.1.

### Immunoblotting

Whole cell extracts from cell lines were generated using radioimmunoprecipitation assay (RIPA) buffer (Thermo Fisher Scientific) supplemented with protease and phosphatase inhibitors (Roche, Basel, Switzerland). Extracted proteins were quantified using Bradford. After separation of 20 to 30 μg extracts on 10% NuPAGE Bis-Tris gels (Thermo Fisher Scientific) under denaturing conditions, proteins were blotted on PVDF membranes under semi-dry conditions using the Trans-Blot® Turbo™ Transfer System (Bio-Rad). Membranes were blocked using the AdvanBlock-Chemi blocking solution. Chemiluminescence was red out on a ImageQuant LAS 4000 (GE Healthcare).

### Antibodies

For immunoblotting, the following antibodies were diluted in AdvanBlock-Chemi blocking solution as indicated and incubated with the membrane for 1h at room temperature or 4°C over night: anti-PHB1 (clone 3F4-2B2; 1:1000; Abnova), anti-PHB2 (clone A-2; 1:1000, Santa Cruz), anti-p53 (clone 7F5 1:1000; Cell Signaling), anti-BTK (clone D3H5; 1:1000 Cell Signaling), anti-p-BTK (Tyr223; 1:1000; Cell Signaling #5082), anti-SYK (clone D3Z1E; 1:1000; Cell signaling), anti-p-SYK (Tyr323; 1:1000; Cell Signaling #2715), anti-mouse IgG HRP coupled (HAF007; 1:5000; R&D Systems), anti-rabbit IgG HRP coupled (HAF008; 1:5000; R&D Systems). For flow cytometry, the following antibodies were diluted 1:200 in PBS: FITC anti-human IgG Fc (clone M1310G05; BioLegend), PE anti-human IgM (clone MHM-88; BioLegend), PE/Cy7 anti-human Ig light chain κ (clone TB28-2; BioLegend), AF647 anti-human Ig light chain λ (RRID: AB_2795758; Southern Biotech).

### qRT-PCR

Total RNA was isolated using the Quick-RNA kit (Zymo Research). After removal of remaining DNA contaminations using the TURBI Dnase Kit (Thermo Fisher), RNA was quantified using a NanoDrop Spectrophotometer (Thermo Fisher). Reverse transcription (RT) of 2.5 µg total RNA was performed in a 20 µl reaction using SuperScript III (Thermo Fisher). After RT and heat inactivation, the sample was diluted to 50 µl with RNase-free water. Target amplification was performed on the CFX96 System (Bio-Rad) using 0.5 µl first-strand cDNA and the SYBR Select Master Mix CFX under the following reaction conditions: initial denaturation/activation for 1 min. at 95°C, denaturation at 95°C for 3 sec, 20 sec of annealing at 54°C, and 20 sec of elongation at 72°C. The following primer combinations were applied: PHB1 (forward: 5’-GAAGGAGTTCACAGAAGCGGT; reverse: 5’-TAGGTGATGTTCCGAGAGCGT), PHB2 (forward: 5’-TTCACTTCAGGATCCCTTGGTT; reverse: 5’-CTGTGATGGCCA CATCATCCA), HPRT1 (forward: 5’-TGACACTGGCAAAACAATGCA; reverse: 5’-GGTCCTTTTCACCAGCAAGCT).

### Panel sequencing

We used 100 ng of DNA for mutational profiling of lymphoma cell lines applying a QIAseq Targeted DNA Custom Panel (Qiagen, Hilden, Germany) with ABC-DLBCL-related target genes (*TNFAIP3, MYD88, CARD11, CD79B, CREBBP, EP300, IRF4, PIM1, TP53, TCF3, PRDM1, TRAF2, TRAF5, MYC, MEF2B, B2M, MAP3K7, TNFRSF11A*, and *ID3*) selected as described in^25^. Quantification and quality control of libraries was conducted using Qubit high-sensitivity double-strand DNA assay kit (Thermo Fisher Scientific) and Agilent 2100 Bioanalyzer (Agilent). Sequencing was performed on the Illumina NextSeq or HiSeq platform with 2 × 151 cycles at an average coverage of 26,500 reads per target region. Variant calling of unique molecular identifier–based sequencing data was performed using smCounter2 as described elsewhere^26^.

### Flow cytometry

For detection of immunoglobulin heavy and light chain surface expression, 1 × 10^6^ target cells were incubated in 500 µl PBS/antibody solution (single stain) for 30 min. at 4°C. After incubation, cells were washed with PBS, resolved in 200 µl PBS and transferred to a 96-well plate. Read out was performed on a CytoFLEX flow cytometer, analysis with FlowJo Software v10.

## Results and Discussion

Analysis of published RNA-seq datasets revealed consistent overexpression of PHB1 and PHB2 transcripts in B cell lymphomas (diffuse large B cell lymphoma, DLBCL^27,28^; chronic lymphocytic leukemia, CLL), compared to healthy controls (Figure 1A). This was mirrored by strong expression in most B cell lymphoma cell lines especially for PHB2 (Figure 1B)^29-31^. We observed a strong correlation between *PHB1* and *PHB2* in healthy blood as well as DLBCL and Burkitt samples, consistent with their known function as heterodimers in the mitochondrial inner membrane in different reanalyzed datasets (Figure 1C and D)^18-21,32^. However, CLL samples showed disproportionally higher *PHB2* levels, resulting in reduced *PHB1/PHB2* correlation, suggesting subtype-specific regulation or functional divergence (Figure 1C). Transcript levels of both types of prohibitins in CLL were independent of IGHV mutation status (Figure 1E) and no strong associations of transcripts with survival were noted (Figure 1F). To explore the association of prohibitins with BCR signaling and to identify potential context-specific roles in this pathway, we assessed *PHB1* and *PHB2* correlations with *BTK* as key mediator of BCR-driven proliferation as well as *SYK* and *RAF1* as important BCR-associated signaling molecules and known interactors of PHB^33,34^. We detected strong positive associations between *PHB1* and *SYK* or *BTK* in Burkitt lymphoma and CLL, and an inverse correlation with *RAF1* (encoding c-RAF) in DLBCL and CLL (Figure 1G). These relationships suggest that PHB1 may be involved in modulating BCR signaling, either by promoting activation through SYK/BTK or by suppressing alternative routes such as RAF1-mediated signaling. In contrast, *PHB2* showed weaker and more variable associations with BCR-associated signaling molecules, with its strongest correlation to *SYK* in Burkitt and a modest inverse correlation to *RAF1* in CLL (Figure 1H).To next assess the therapeutic potential of prohibitin modulation in B cell malignancies, we evaluated the impact of the five PHB-targeting molecules FL3, Mel6, Mel56, IN44, and Fluorizoline (Figure 2A) on a range of lymphoma cell models. While the MEC-1 M-CLL line served as our primary model alongside non-malignant M2-10B4 stromal cells (Figure 2), we also tested additional B and T cell lymphoma lines (Figure 3). In MEC-1 cells, FL3 and Mel56 led to a depletion of PHB1/2 proteins, whereas the other modulators had no or minimal effects (Figure 2B). Transcriptional analysis suggested that these protein-level changes were only partially due to altered gene expression (Figure 2C). FL3, and to some extent also Fluorizoline, triggered upregulation of *PHB* transcripts, likely reflecting a feedback or stress response to counteract protein destabilization, while Mel56 selectively suppressed *PHB1* transcripts in MEC-1 cells (Figure 2C). Functionally, FL3 and Mel56 were the most potent inhibitors of proliferation and viability in MEC-1 cells, whereas Mel6 primarily slowed proliferation without inducing significant cytotoxicity. IN44 and Fluorizoline had minimal impact on MEC-1 growth (Figure 2D). Importantly, none of the compounds reduced viability in non-malignant cells, though FL3 suppressed their proliferation

**Figure 1.**
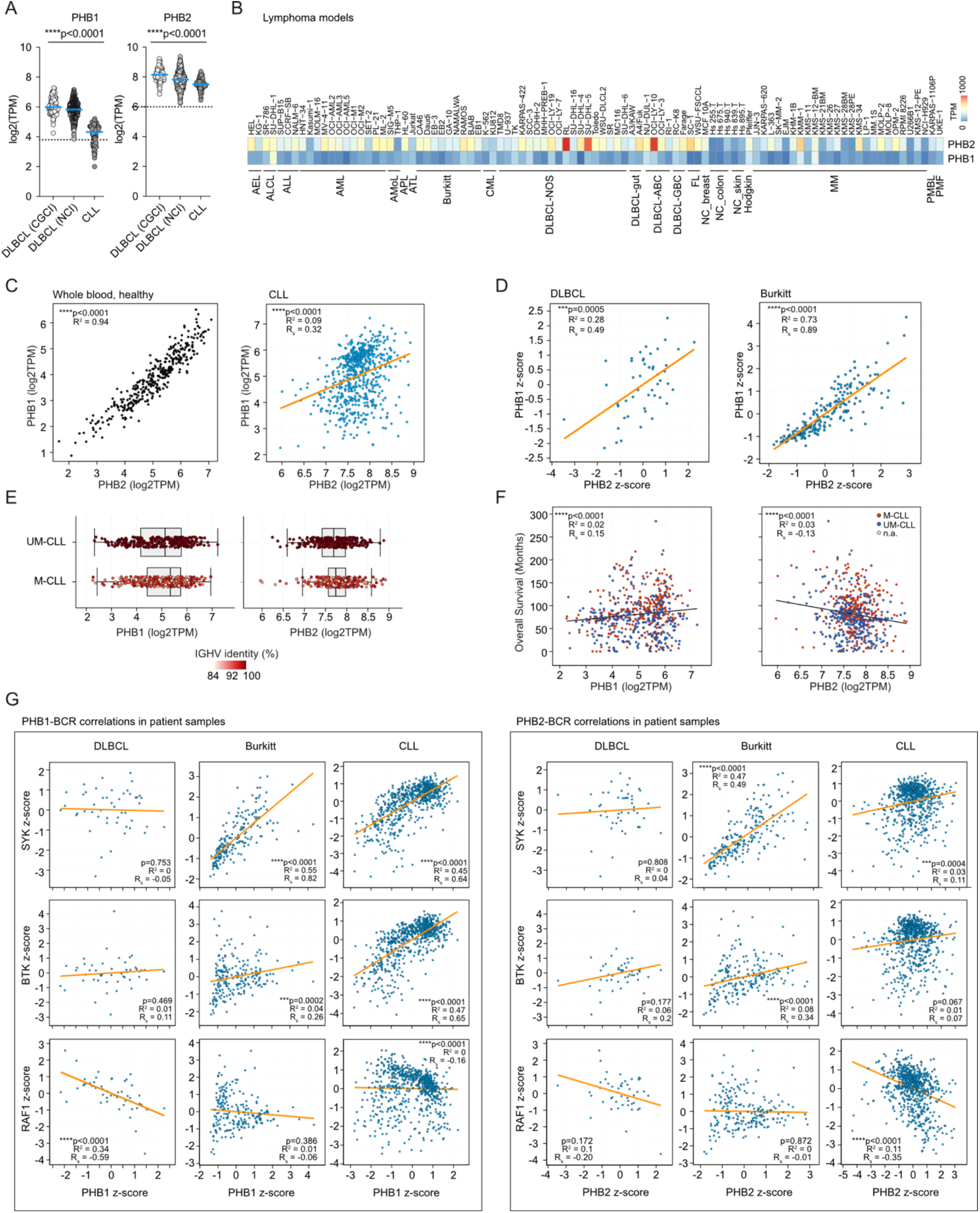
Prohibitin expression in B cell neoplasias. **(A)** Mean *PHB1* and *PHB2* expression in two different DLBCL cohorts (DLBCL (NCI)^28^, n=481; DLBCL (CGCI)^27^, n=98) and a CLL^35^ cohort (n=69). RNAseq data accessed via UCSC Xena. Dotted line represents mean *PHB1* or *PHB2* expression in healthy blood (TCGA data). Statistics: Ordinary one-way ANOVA. **(B)** Expression of *PHB1* and *PHB2* in different B cell lymphoma models. **(C)** Spearman correlation of *PHB1*/*PHB2* expression in RNAseq data derived from healthy blood accessed via GEPIA2 and CLL from Knisbacher et al.^21^ accessed via cBioportal. **(D)** Spearman correlation of *PHB1*/*PHB2* expression in DLBCL and Burkitt lymphoma samples (both from Thomas et al.^20^) accessed via cBioportal. **(E)** Expression levels of *PHB1*/*PHB2* depending on IGHV mutation status in CLL. Samples from Knisbacher et al.^21^ accessed via cBioportal. **(F)** Spearman correlation of Overall Survival with *PHB1*/*PHB2* expression depending on IGHV mutation status in CLL. Samples from Knisbacher et al.^21^ accessed via cBioportal. **(G)** Spearman correlation of *PHB1* and *PHB2* expression with BCR-related signaling molecules in DLBCL, Burkitt, and CLL samples from (C) and (D). Correlation coefficient *R*^2^, Spearman correlation coefficients (*r*_S_), and *p* values are indicated in all respective panels.

**Figure 2.**
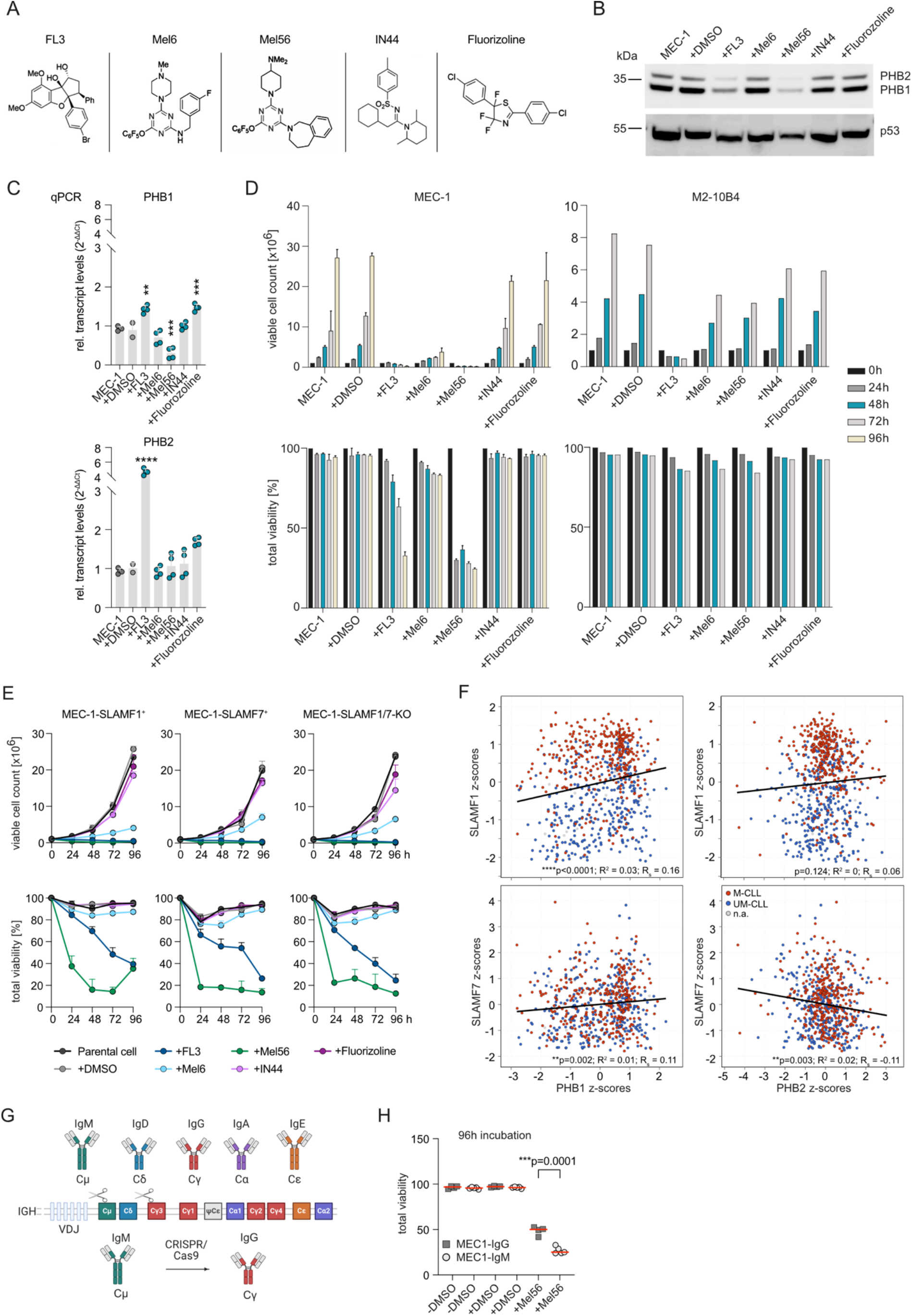
Effects of PHB modulation on PHB protein abundancies, transcript levels, proliferation and viability in a CLL cell line model. **(A)** PHB-targeting small molecules tested in this study. **(B)** PHB1 and PHB2 protein abundancies after 48h after PHB targeting. **(C)** Transcript levels of PHB1/2 after 48h PHB targeting. **(D)** Proliferation and viability of MEC-1 CLL and M2-10B4 stromal cells over time after PHB targeting. n=3 for all. **(E)** Proliferation and viability of MEC-1 CLL deficient for SLAMF1/7 or overexpressing either SLAMF1 or SLAMF7 after treatment with PHB-targeting small molecules. n=3 for all. **(F)** Spearman correlation of *SLAMF1, SLAMF7, PHB1* and *PHB2* expression in primary CLL samples dependent on IGHV status. Samples from Knisbacher et al.^21^ accessed via cBioportal. Correlation coefficient *R*^2^, Spearman correlation coefficients (*r*_S_), and *p* values are indicated in all respective panels. **(G)** Schematic representation of CRISPR-Cas9-mediated switch of IgM to IgG. **(H)** Effect of Mel56 on cell viability depending on BCR IgM versus IgG class. n=3.

**Figure 3.**
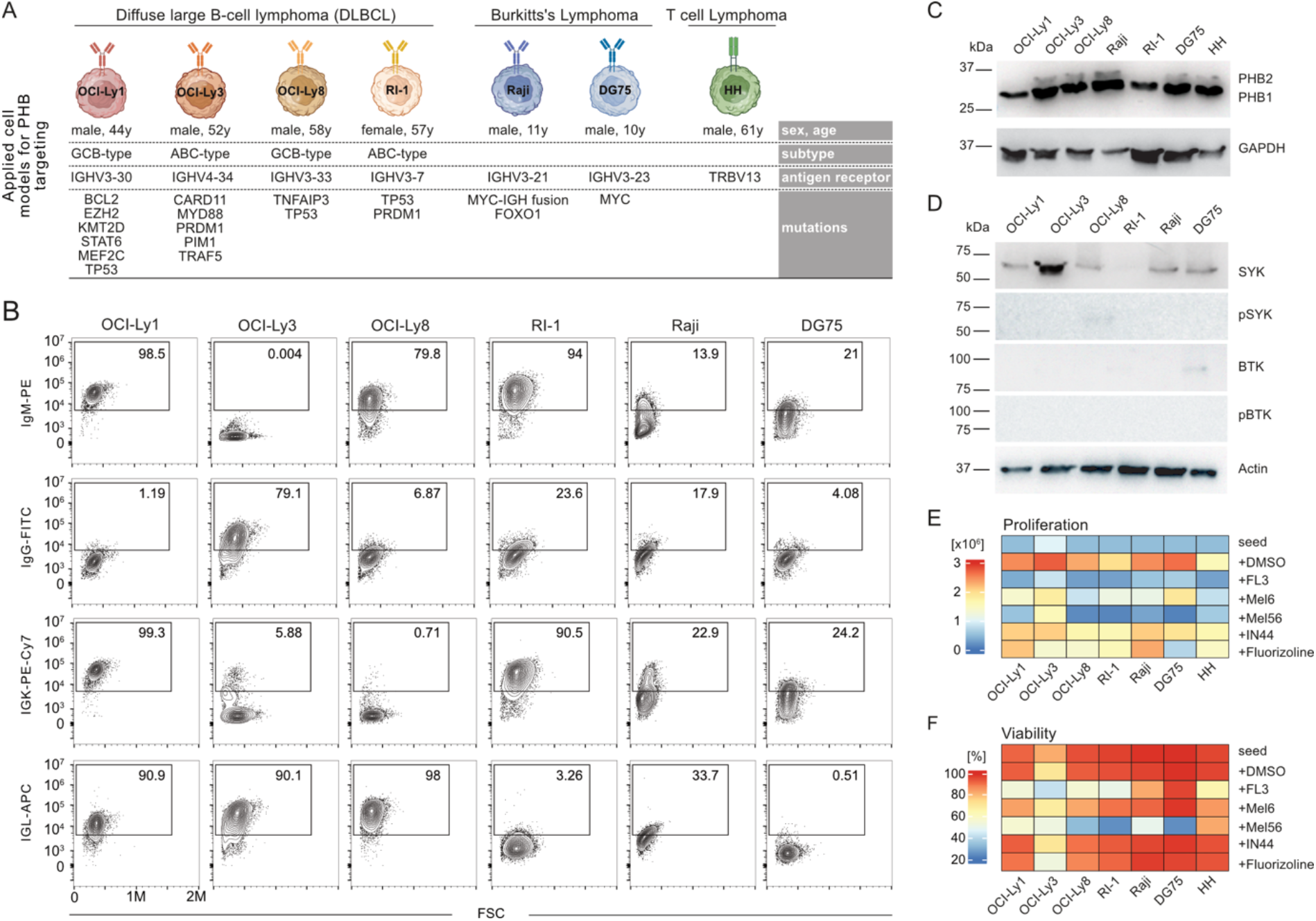
PHB modulation in non-CLL lymphoma models. **(A)** B and T lymphoma cell lines used in this study. **(B)** Classification of surface Ig expression in applied B cell models using flow cytometry. **(C)** Detection of PHB1 and PHB2 protein abundancies in the applied lymphoma cells using immunoblotting. **(D)** Protein abundancies and activation of proximal BCR kinases in the applied B cell models as detected by immunoblotting. **(E)** Proliferation of cell models after treatment with prohibitin-targeting small molecules (n=3). Heatmap represents total cell number after 96h treatment. Seed-number included for comparison. **(F)** Cell viability of lymphoma models after 96h treatment with prohibitin-targeting small molecules displayed as heatmap (n=3).

(Figure 2D). SLAMF1 and SLAMF7 have been identified as PHB-interacting proteins and increased surface expression for SLAMF1/7 has been described to outcompete the BCR for PHB binding resulting in reduced BCR signaling and better prognosis in CLL^13^. However, when testing the PHB-targeting small molecules in MEC-1 cells deficient for SLAMF1/7 or overexpressing either SLAMF1 or SLAMF7, we did not detect specific shifts in drug response, but rather recapitulated proliferation and viability patterns as observed for the parental MEC-1 cell line (Figure 2E). Bulk CLL datasets nevertheless showed weak yet significant positive correlations between *SLAMF1* and *PHB1* and *PHB2*, especially in M-CLL, while *SLAMF7* correlated positively with *PHB1* but inversely with PHB2 independent of IGHV status (Figure 2F). Rather than invalidating the functional role of the SLAMF-PHB axis, these results highlight the pleiotropic roles of PHBs and the layered biology of CLL, where the reported FL3-mediated block of MYC translation by PHB-eIF4F in CLL^31^ or the FL3-or Mel56-mediated block of the c-RAF1-ERK axis^29,36^ may represent more dominant functional consequences that mask the additive (SLAMF overexpression) or opposing (SLAMF deficiency) effects of the SLAMF-PHB axis on proliferation. In line with this notion, we observed that MEC-1 cells engineered to express an IgG-type BCR (Figure 2G) were less sensitive to Mel56 as their counterparts expressing an IgM-type BCR which is the predominant configuration for BCR-PHB interaction (Figure 2H).

To explore lymphoma entity-specific drug responses, we next profiled the PHB-targeting small molecules in a set of non-Hodgkin lymphoma cell lines (DLBCL; Burkitt lymphoma) as well as a T cell lymphoma model (HH) (Figure 3A). We determined all antigen receptor configurations by BCR/TCR immunosequencing, performed gene panel sequencing to profile the B lymphomas for common driver mutations (Figure 3A), and also analyzed all B lymphoma cells for their BCR type and expression level by flow cytometry to detect potential context-dependent roles of prohibitins (Figure 3B). Immunoblotting revealed PHB1 expression in all cell lines, while PHB2 signals in DLBCL cell lines OCI-Ly1 and RI-1 were not detectable (Figure 3C). Although SYK expression was more pronounced than BTK, we did not observe activation of either kinase in resting cells (Figure 3D). The patterns of antiproliferative and cytotoxic effects were comparable to the MEC-1 CLL model with FL3 and Mel56 again being most antiproliferative and inducing robust cytotoxicity (Figure 3E and F). Of note, the IgG positive line OCI-Ly3 was relatively resistant to Mel56 (Figure 3E and F). This cell line also exhibits a clear mutational profile for constitutive NF-κB activation which may counteract PHB targeting. Interestingly, the T cell lymphoma cell line HH was virtually unaffected by PHB targeting except for FL3 that overall showed lowest selectivity for B cell lines given also its effect on stromal cells described above. Notably, Fluorizoline selectively reduced proliferation but not viability in the Burkitt model DG75, while not affecting any other tested cell line (Figure 3E and F). These data highlighted substance-specific vulnerabilities and potential safety in non-B cell contexts.

Taken together, our data raise the possibility that PHB-targeting small molecules could exert selective effects in B cell malignancies and warrant further investigation. The pleiotropic functions of PHBs make them also potential candidates in cases of resistance to established BCR pathway inhibitors. However, the exact targeting mechanisms for the here tested compounds as well as the context-dependent dominant functions of PHBs in different lymphomas and their immunomodulatory potential require further investigation. This may also help to identify settings where combinatorial approaches with SYK or BTK inhibitors may prove synergistic.

## Acknowledgments

We thank Yiqing Du for excellent technical assistance. This project was funded by the DJCLS.

## Author contributions

Idea & design of research project: MB, CS. Experimental work: CS, MK. Supply of material: LD. Data analysis: CS, MB. Drafting manuscript: CS, MB.

## Competing interests

The authors declare that they have no competing interests.

## Notes

### Competing Interest Statement

The authors have declared no competing interest.

## References

1. Kuppers, R. (2005). Mechanisms of B-cell lymphoma pathogenesis. Nat Rev Cancer 5, 251–262. 10.1038/nrc1589.

2. Silkenstedt, E., Salles, G., Campo, E., and Dreyling, M. (2024). B-cell non-Hodgkin lymphomas. Lancet 403, 1791–1807. 10.1016/S0140-6736(23)02705-8.

3. Kipps, T.J., Stevenson, F.K., Wu, C.J., Croce, C.M., Packham, G., Wierda, W.G., O’Brien, S., Gribben, J., and Rai, K. (2017). Chronic lymphocytic leukaemia. Nat Rev Dis Primers 3, 16096. 10.1038/nrdp.2016.96.

4. Young, R.M., Phelan, J.D., Wilson, W.H., and Staudt, L.M. (2019). Pathogenic B-cell receptor signaling in lymphoid malignancies: New insights to improve treatment. Immunol Rev 291, 190–213. 10.1111/imr.12792.

5. Sun, R.F., Yu, Q.Q., and Young, K.H. (2018). Critically dysregulated signaling pathways and clinical utility of the pathway biomarkers in lymphoid malignancies. Chronic Dis Transl Med 4, 29–44. 10.1016/j.cdtm.2018.02.001.

6. Shaffer, A.L., 3rd, Young, R.M., and Staudt, L.M. (2012). Pathogenesis of human B cell lymphomas. Annu Rev Immunol 30, 565–610. 10.1146/annurev-immunol-020711-075027.

7. Clara, J.A., Monge, C., Yang, Y., and Takebe, N. (2020). Targeting signalling pathways and the immune microenvironment of cancer stem cells -a clinical update. Nat Rev Clin Oncol 17, 204–232. 10.1038/s41571-019-0293-2.

8. Rajalingam, K., Wunder, C., Brinkmann, V., Churin, Y., Hekman, M., Sievers, C., Rapp, U.R., and Rudel, T. (2005). Prohibitin is required for Ras-induced Raf-MEK-ERK activation and epithelial cell migration. Nat Cell Biol 7, 837–843. 10.1038/ncb1283.

9. Wei, Y., Chiang, W.C., Sumpter, R., Jr., Mishra, P., and Levine, B. (2017). Prohibitin 2 Is an Inner Mitochondrial Membrane Mitophagy Receptor. Cell 168, 224–238 e210. 10.1016/j.cell.2016.11.042.

10. Yan, C., Gong, L., Chen, L., Xu, M., Abou-Hamdan, H., Tang, M., Desaubry, L., and Song, Z. (2020). PHB2 (prohibitin 2) promotes PINK1-PRKN/Parkin-dependent mitophagy by the PARL-PGAM5-PINK1 axis. Autophagy 16, 419–434. 10.1080/15548627.2019.1628520.

11. Kasashima, K., Ohta, E., Kagawa, Y., and Endo, H. (2006). Mitochondrial functions and estrogen receptor-dependent nuclear translocation of pleiotropic human prohibitin 2. J Biol Chem 281, 36401–36410. 10.1074/jbc.M605260200.

12. Fusaro, G., Dasgupta, P., Rastogi, S., Joshi, B., and Chellappan, S. (2003). Prohibitin induces the transcriptional activity of p53 and is exported from the nucleus upon apoptotic signaling. J Biol Chem 278, 47853–47861. 10.1074/jbc.M305171200.

13. von Wenserski, L., Schultheiss, C., Bolz, S., Schliffke, S., Simnica, D., Willscher, E., Gerull, H., Wolters-Eisfeld, G., Riecken, K., Fehse, B., et al. (2021). SLAMF receptors negatively regulate B cell receptor signaling in chronic lymphocytic leukemia via recruitment of prohibitin-2. Leukemia 35, 1073–1086. 10.1038/s41375-020-01025-z.

14. Moreno, P., Fexova, S., George, N., Manning, J.R., Miao, Z., Mohammed, S., Munoz-Pomer, A., Fullgrabe, A., Bi, Y., Bush, N., et al. (2022). Expression Atlas update: gene and protein expression in multiple species. Nucleic Acids Res 50, D129–D140. 10.1093/nar/gkab1030.

15. Gao, J., Aksoy, B.A., Dogrusoz, U., Dresdner, G., Gross, B., Sumer, S.O., Sun, Y., Jacobsen, A., Sinha, R., Larsson, E., et al. (2013). Integrative analysis of complex cancer genomics and clinical profiles using the cBioPortal. Sci Signal 6, pl1. 10.1126/scisignal.2004088.

16. Goldman, M.J., Craft, B., Hastie, M., Repecka, K., McDade, F., Kamath, A., Banerjee, A., Luo, Y., Rogers, D., Brooks, A.N., et al. (2020). Visualizing and interpreting cancer genomics data via the Xena platform. Nat Biotechnol 38, 675–678. 10.1038/s41587-020-0546-8.

17. Tang, Z., Kang, B., Li, C., Chen, T., and Zhang, Z. (2019). GEPIA2: an enhanced web server for large-scale expression profiling and interactive analysis. Nucleic Acids Res 47, W556–W560. 10.1093/nar/gkz430.

18. Gao, G.F., Parker, J.S., Reynolds, S.M., Silva, T.C., Wang, L.B., Zhou, W., Akbani, R., Bailey, M., Balu, S., Berman, B.P., et al. (2019). Before and After: Comparison of Legacy and Harmonized TCGA Genomic Data Commons’ Data. Cell Syst 9, 24–34 e10. 10.1016/j.cels.2019.06.006.

19. Klijn, C., Durinck, S., Stawiski, E.W., Haverty, P.M., Jiang, Z., Liu, H., Degenhardt, J., Mayba, O., Gnad, F., Liu, J., et al. (2015). A comprehensive transcriptional portrait of human cancer cell lines. Nat Biotechnol 33, 306–312. 10.1038/nbt.3080.

20. Thomas, N., Dreval, K., Gerhard, D.S., Hilton, L.K., Abramson, J.S., Ambinder, R.F., Barta, S., Bartlett, N.L., Bethony, J., Bhatia, K., et al. (2023). Genetic subgroups inform on pathobiology in adult and pediatric Burkitt lymphoma. Blood 141, 904–916. 10.1182/blood.2022016534.

21. Knisbacher, B.A., Lin, Z., Hahn, C.K., Nadeu, F., Duran-Ferrer, M., Stevenson, K.E., Tausch, E., Delgado, J., Barbera-Mourelle, A., Taylor-Weiner, A., et al. (2022). Molecular map of chronic lymphocytic leukemia and its impact on outcome. Nat Genet 54, 1664–1674. 10.1038/s41588-022-01140-w.

22. Weber, K., Bartsch, U., Stocking, C., and Fehse, B. (2008). A multicolor panel of novel lentiviral “gene ontology” (LeGO) vectors for functional gene analysis. Mol Ther 16, 698–706. 10.1038/mt.2008.6.

23. Tintelnot, J., Metz, S., Trentmann, M., Oberle, A., von Wenserski, L., Schultheiss, C., Braig, F., Kriegs, M., Fehse, B., Riecken, K., et al. (2019). Cancer Cells Expressing Oncogenic Rat Sarcoma Show Drug-Addiction Toward Epidermal Growth Factor Receptor Antibodies Mediated by Sustained MAPK Signaling. Front Oncol 9, 1559. 10.3389/fonc.2019.01559.

24. Cheong, T.C., Compagno, M., and Chiarle, R. (2016). Editing of mouse and human immunoglobulin genes by CRISPR-Cas9 system. Nat Commun 7, 10934. 10.1038/ncomms10934.

25. Schultheiss, C., Paschold, L., Mohebiany, A.N., Escher, M., Kattimani, Y.M., Muller, M., Schmidt-Barbo, P., Mensa-Vilaro, A., Arostegui, J.I., Boursier, G., et al. (2024). A20 haploinsufficiency disturbs immune homeostasis and drives the transformation of lymphocytes with permissive antigen receptors. Sci Adv 10, eadl3975. 10.1126/sciadv.adl3975.

26. Xu, C., Gu, X., Padmanabhan, R., Wu, Z., Peng, Q., DiCarlo, J., and Wang, Y. (2019). smCounter2: an accurate low-frequency variant caller for targeted sequencing data with unique molecular identifiers. Bioinformatics 35, 1299–1309. 10.1093/bioinformatics/bty790.

27. Heath, A.P., Ferretti, V., Agrawal, S., An, M., Angelakos, J.C., Arya, R., Bajari, R., Baqar, B., Barnowski, J.H.B., Burt, J., et al. (2021). The NCI Genomic Data Commons. Nat Genet 53, 257–262. 10.1038/s41588-021-00791-5.

28. Phelan, J.D., Young, R.M., Webster, D.E., Roulland, S., Wright, G.W., Kasbekar, M., Shaffer, A.L., 3rd, Ceribelli, M., Wang, J.Q., Schmitz, R., et al. (2018). A multiprotein supercomplex controlling oncogenic signalling in lymphoma. Nature 560, 387–391. 10.1038/s41586-018-0290-0.

29. Bentayeb, H., Aitamer, M., Petit, B., Dubanet, L., Elderwish, S., Desaubry, L., de Gramont, A., Raymond, E., Olivrie, A., Abraham, J., et al. (2019). Prohibitin (PHB) expression is associated with aggressiveness in DLBCL and flavagline-mediated inhibition of cytoplasmic PHB functions induces anti-tumor effects. J Exp Clin Cancer Res 38, 450. 10.1186/s13046-019-1440-4.

30. Wierz, M., Pierson, S., Chouha, N., Desaubry, L., Francois, J.H., Berchem, G., Paggetti, J., and Moussay, E. (2018). The prohibitin-binding compound fluorizoline induces apoptosis in chronic lymphocytic leukemia cells ex vivo but fails to prevent leukemia development in a murine model. Haematologica 103, e154–e157. 10.3324/haematol.2017.175349.

31. Largeot, A., Klapp, V., Viry, E., Gonder, S., Fernandez Botana, I., Blomme, A., Benzarti, M., Pierson, S., Duculty, C., Marttila, P., et al. (2023). Inhibition of MYC translation through targeting of the newly identified PHB-eIF4F complex as a therapeutic strategy in CLL. Blood 141, 3166–3183. 10.1182/blood.2022017839.

32. Lange, F., Ratz, M., Dohrke, J.N., Le Vasseur, M., Wenzel, D., Ilgen, P., Riedel, D., and Jakobs, S. (2025). In situ architecture of the human prohibitin complex. Nat Cell Biol 27, 633–640. 10.1038/s41556-025-01620-1.

33. Paris, L.L., Hu, J., Galan, J., Ong, S.S., Martin, V.A., Ma, H., Tao, W.A., Harrison, M.L., and Geahlen, R.L. (2010). Regulation of Syk by phosphorylation on serine in the linker insert. J Biol Chem 285, 39844–39854. 10.1074/jbc.M110.164509.

34. Ande, S.R., Xu, Y.X.Z., and Mishra, S. (2017). Prohibitin: a potential therapeutic target in tyrosine kinase signaling. Signal Transduct Target Ther 2, 17059. 10.1038/sigtrans.2017.59.

35. Consortium, I.T.P.-C.A.o.W.G. (2020). Pan-cancer analysis of whole genomes. Nature 578, 82–93. 10.1038/s41586-020-1969-6.

36. Najem, A., Krayem, M., Sabbah, S., Pesetti, M., Journe, F., Awada, A., Desaubry, L., and Ghanem, G.E. (2023). Targeting Prohibitins to Inhibit Melanoma Growth and Overcome Resistance to Targeted Therapies. Cells 12. 10.3390/cells12141855.

